# Distinct co-expression networks using multi-omic data reveal novel interventional targets in HPV-positive and negative head-and-neck squamous cell cancer

**DOI:** 10.1101/236133

**Authors:** Raquel L. Costa, Mariana Boroni, Marcelo A. Soares

## Abstract

The human papillomavirus (HPV) is present in a significant fraction of head-and-neck squamous cell cancer (HNSCC). However, a comprehensive understanding of disease progression profiles comparing HPV+ and HPV- HNSCC cases is still lacking. The main goal of this study was to identify distinct co-expression patterns between HPV+ and HPV- HNSCC and to provide insights into potential regulatory mechanisms/effects (such as methylation and mutation) within the analyzed networks. For conducting this, we selected 276 samples from The Cancer Genome Atlas database comprising data of gene expression, methylation profiles and mutational patterns, in addition to clinical information (HPV status and tumor staging). We further added external information such as the identification of transcription factors to the networks. Genes were selected as differentially expressed and differentially methylated based on HPV status, of which 12 genes were doubly selected, including *SYCP2, GJB6, FLRT3, PITX2* and *CCNA1*. Weight correlation network analysis was used to identify co-expression modules and a systematic approach was applied to refine them and identify key regulatory elements integrating results from the other omics. Three main modules were associated with distinct co-expression patterns in HPV+ versus HPV- HNSCC. The molecular signatures found were mainly related to cell fate specification, keratinocyte differentiation, focal adhesion and regulation of protein oligomerization. This study provides comprehensive insights into complex genetic and epigenetic particularities in the development and progression of HNSCC in patients according to HPV status, identifying unseen gene interactions, and may contribute to unveiling specific genes/pathways as novel therapeutic targets for HNSCC.

## Background

Head-and-neck squamous cell carcinoma (HNSCC) is a heterogeneous malignancy which accounts for approximately 300,000 deaths each year worldwide (1, 2). Smoking, alcohol, and infections by high-risk human papillomavirus (HPV) are among the main risk factors for the development of the disease. The incidence of HPV- associated HNSCC is around 25% of the reported cases worldwide, with an even higher proportion of oropharyngeal cancer, and a predominance of infection by HPV- 16 and HPV- 18 types among those cases (3,4).

The development and progression of HNSCC occur by molecular deregulation events in many levels, including the accumulation of somatic mutations and changes in methylation profiles. Both those events result in differences in gene expression levels and downstream signaling pathways. In general, patients diagnosed with HNSCC and HPV+ have a better prognosis (regardless of the treatment strategies) compared with the patients without HPV (HPV-) in the same anatomical site (5–7). Although the molecular mechanisms involved in those difference are not fully understood, mutations of in the *TP53* gene are massively more detected in HPV- compared to HPV+ tumors (8–10).

With the advancement of high-throughput technologies, such as next-generation sequencing (NGS), efforts have been made to identify molecular characteristics that differentiate the profiles of HPV+ and HPV- HNSCC. Studies involving gene expression profiles have identified potential marker genes within each context. Masterson and colleagues (2015) (11) identified markers of early-stage HPV+ oropharyngeal squamous cell carcinomas. Wood et al. (2016) (12) identified distinct immune signatures in tumor-infiltrating lymphocytes (TILs), more specifically in B-cells, related to the adaptive immune response against HPV in those tumors. Gene expression involving microarray technology in HPV+ versus HPV- HNSCC has also been studied (13, 14). Other studies considered differences in methylation profiles. Esposti et al. (2017) (15), for example, identified novel epigenetic signatures of HPV infection in HNSCC independent of the anatomical site. Studies involved more than one omic are increasing in the recent literature. Seiwert et al. (2015) (16) used mutation and copy-number variation data to find unique mutations and aberrations in HPV+ HNSCC. Characterization of HNSCC subgroups using copy number alteration and transcriptome data were used in some studies (17, 18). The Cancer Genome Atlas (TCGA) consortium conducted a large study containing multi-platform and different types of tumors, including HNSCC. In 2015 the consortium carried out a comprehensive characterization of HNSCC samples including the identification of their HPV status (10).

Although several studies have already been described for HPV+ and HPV- HNSCC, the incorporation of multi-omic data in the analysis is a recent approach and it is still scarcely explored (19–22), mainly in interaction networks. In gene interaction networks, such integration is essential in the construction and functional understanding of the connections between genes at multiple levels (23). With advances in research such as the TCGA mentioned above and other multiomic repositories, it becomes possible to analyze a diversity of tumors through different platforms and technologies (24). In the present study, we have used HNSCC multi-omic data from the TCGA to explore the differences between gene coexpression networks of HPV+ and HPV- disease profiles. We first collected genes with significant differences in promoter methylation and gene expression profiles for each stage of the disease (Differentially Methylated Genes – DMG – and Differentially Expressed Genes – DEG –, respectively). The intersection among DMG and DEG showed the negative correlations between the levels of methylation and expression, suggesting that these genes have their expression levels regulated by methylation alteration patterns in their promoter. Based on global gene expression patterns, we applied Weighted Correlation Network Analysis (WGCNA) to identify gene modules associated with HPV status, followed by a computational strategy pipeline designed by us to refine the modules and build the networks for specific HPV profiles. In our results, the networks significantly associated with HPV statuses showed different connection patterns and brought new insights into mechanisms associated with HPV+ HNSCC. To our knowledge, this is the first study to conduct a gene network reconstruction via the integration of multi-omic sets for HPV+ and HPV- HNSCC.

## Results

### Gene expression profiles are influenced by methylation status in HPV+ and HPV- HNSCC

The datasets studied were preprocessed and analyzed using the flowchart represented in Figure 1. The preprocessing TCGA dataset for RNA-Seq level-3 resulted in 20,502 analyzed genes. For DNA methylation level-3, the dataset resulted in 14,861 analyzed genes. Two hundred and twenty-three DEG and 359 DMG were selected when comparing HPV+ and HPV- tumor samples. For methylation, only probes corresponding to the TSS200 annotation, following the strategy described in the ‘Methods’ were considered. The genes were selected using the limma package (25) with restrictive parameters (False Discovery Rate, *FDR* < 0.01, absolute (abs) Fold Change, *FC* > 3 and abs *FC* > 4 for expression and methylation levels, respectively) and evaluated for differences of HNSCC with HPV+ versus HPV- profiles within each disease stage (I – IV; Figure 2A).

**Fig. 1.**
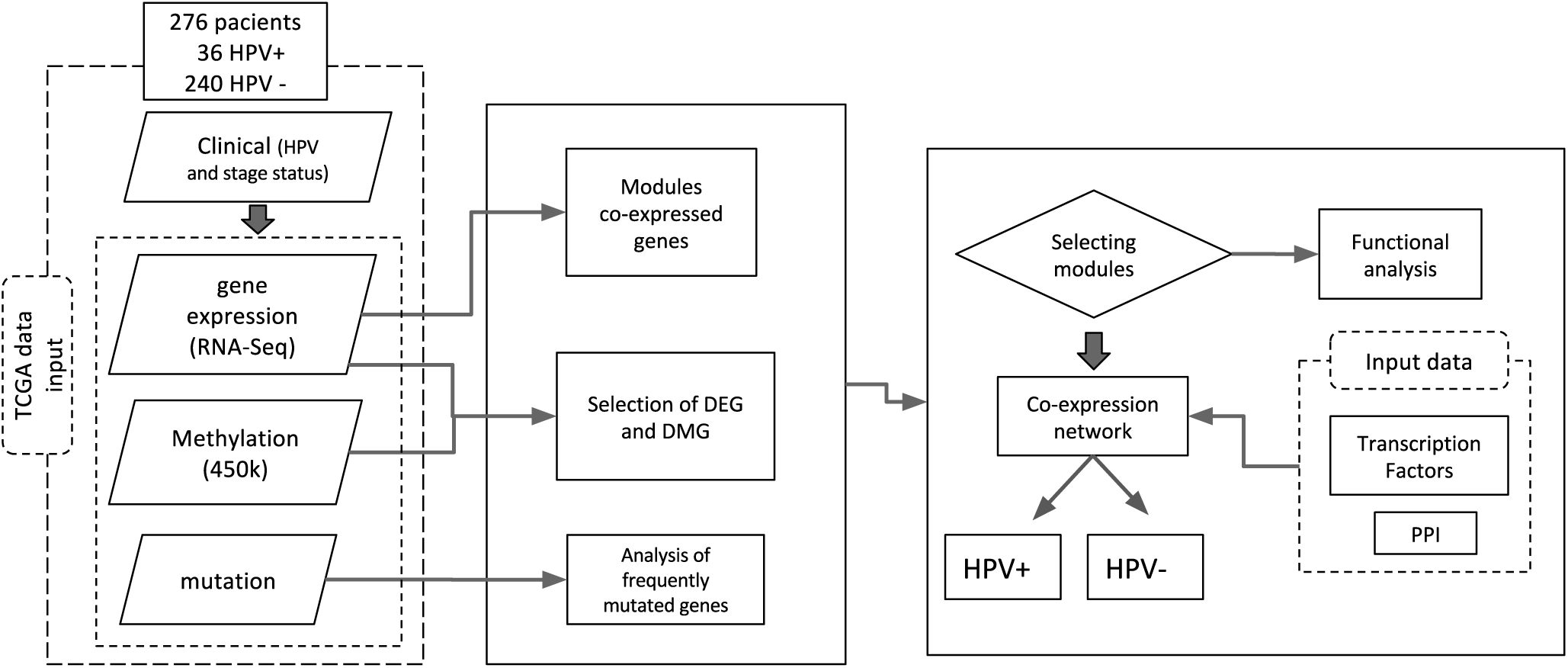
Flow diagram of the methodology applied to this study. The representation includes dataset preparation (left panel, dashed boxes), processes and analysis (middle and right panels, solid boxes).

**Fig. 2.**
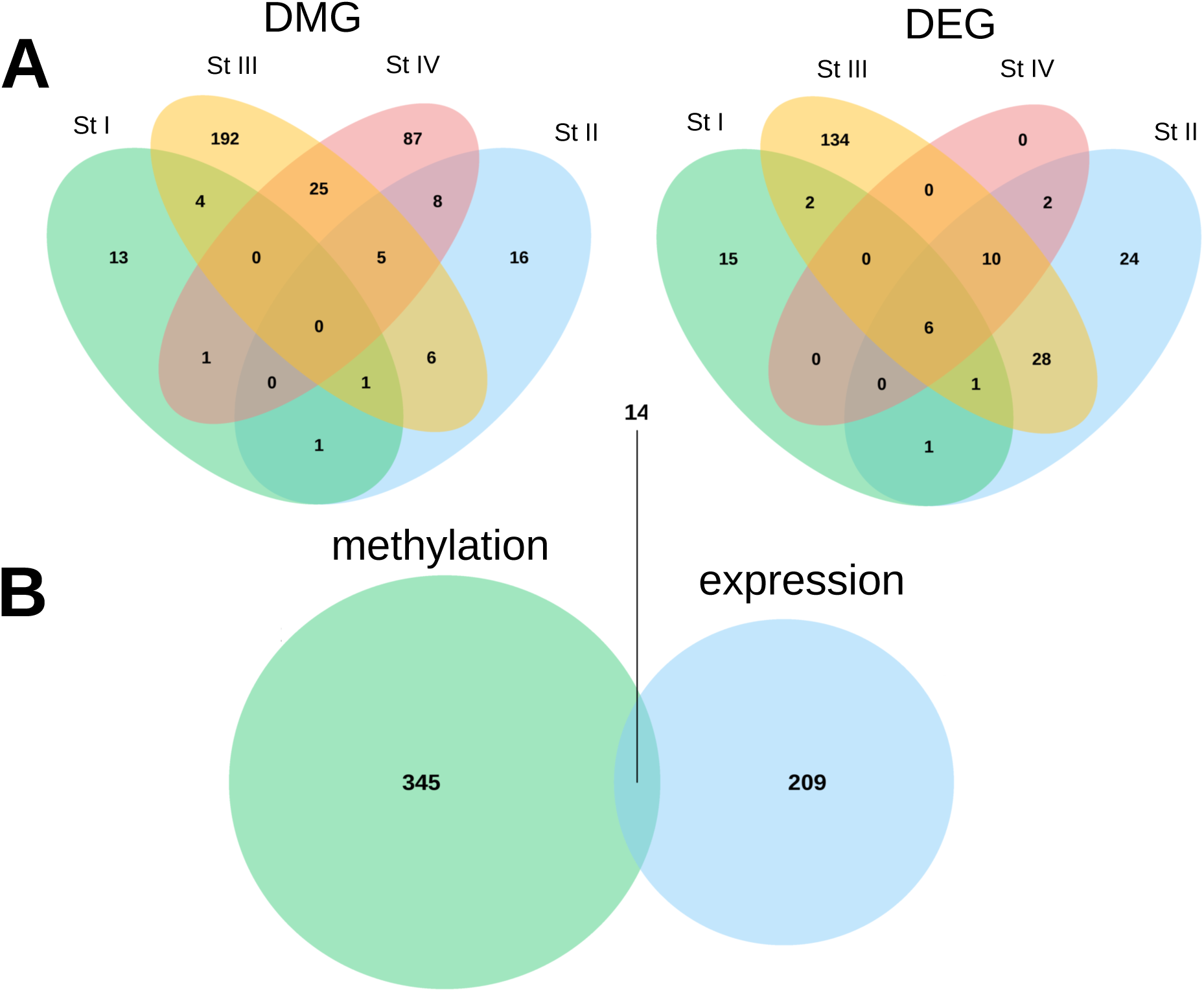
Venn diagrams showing the number of selected genes with differential methylation (DMG; left panels) and expression (DEG; right panels) profiles between HPV+ and HPV-tumor samples. In **A,** DEG and DMG are stratified by tumor stages (st I to IV, represented by distinct colors), respectively. In **B,** the overlap of genes selected in **A** by differential methylation and expression among HPV+ versus HPV- is shown.

The overlapping between DMG and DEG resulted in 14 genes which were doubly differential (Figure 2B). For this selection, a Pearson’s correlation (PC) was carried out between the expression and the methylation values (Figure 3A-L). The *PNLDC1* and *CNTN1* genes were excluded from subsequent analysis due to their similar correlation profiles in both HPV+ and HPV- samples and both were differentially selected only in early stages, for which a limited number of samples were available for analysis. For all 12 genes evaluated, we found PC values < 0, showing a negative correlation between the two parameters. For seven genes (58,3%), the PC obtained for HPV+ HNSCC were higher than for HPV. Our results are consistent with the knowledge of methylated promoter regions negatively regulating gene expression levels. In HPV+ cases, the *SYCP2, MEI1, UGT8, ZFR2* and *SOX30* genes were overexpressed when compared to the HPV- cases, an observation that was coupled with a decreased promoter methylation profile in the former (Figures 3A-E). Conversely, the *FLRT3, PITX2* and *SPRR2G* genes were underexpressed in HPV+ cases compared to the HPV- cases (Figures 3G-I). In those cases, a stronger negative correlation was seen in the HPV+ cases. On the other hand, the *GJB6* gene also exhibited underexpression in HPV+ cases, but a stronger negative correlation (rho ≤ 0.5) in the HPV- cases. As expected, for those four genes, a consistent stronger promoter methylation was observed in the HPV+ cases (Figures 3G-J).

**Fig. 3.**
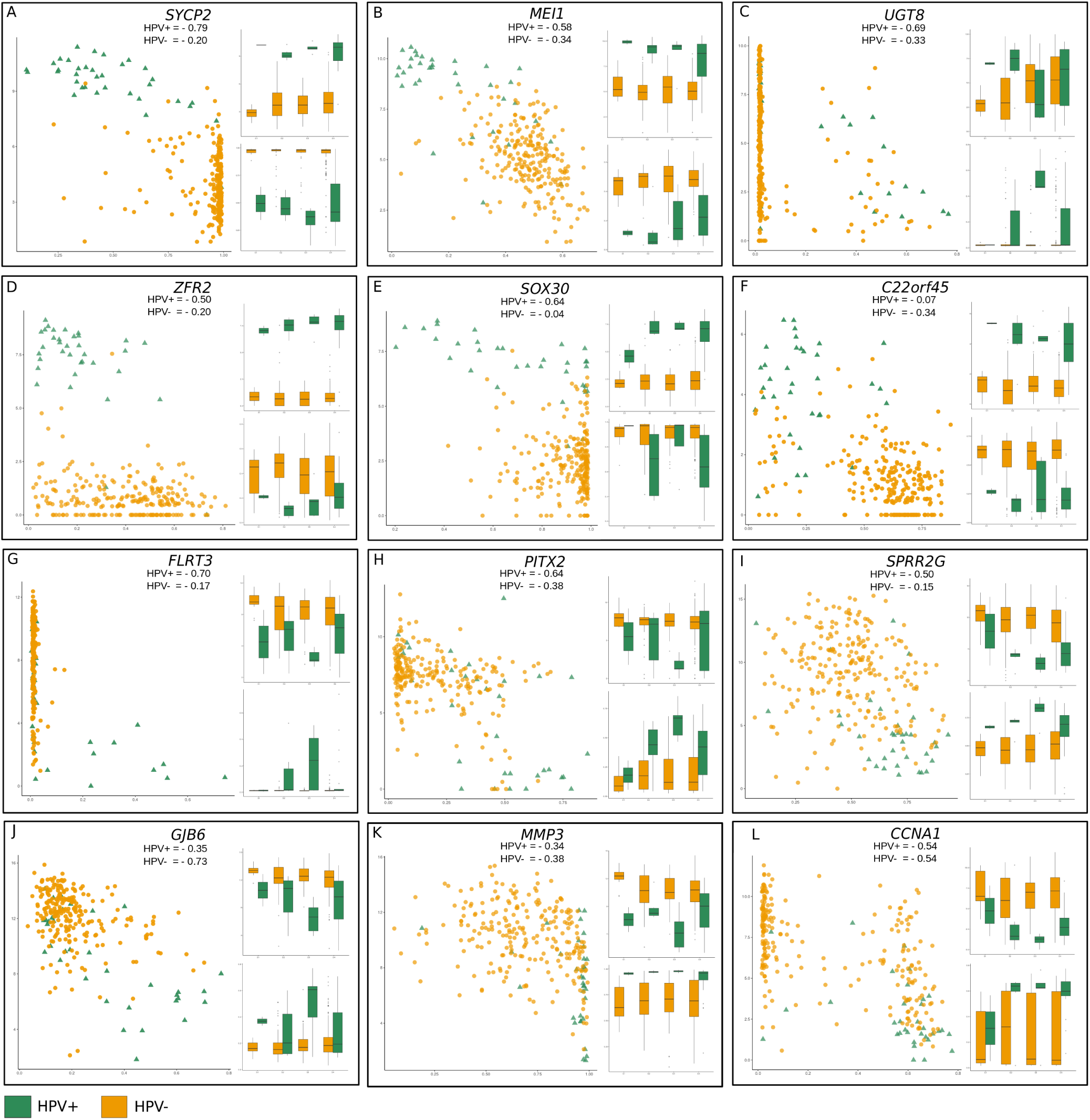
Negative correlation between gene expression and promoter methylation levels of genes doubly selected (**A-L**). For each gene, a scatter plot shows the correlation among methylation (x-axis) and expression levels (y-axis) for each profile (yellow circles for HPV- and green triangles for HPV+ samples). In each inset, the expression (upper panels) and the methylation levels (lower panels) are compared for each tumor stage (I to IV), using the same color codes for HPV+ and HPV- statuses.

### Gene modules were significantly associated with HPV status

Gene co-expression networks were constructed using the WGCNA approach. The method calculates correlations among genes across the samples and applies a power function to determine the connection strengths between genes resulting in a scale-free network (26, 27). Due to computational time, we used the 8,000 most variant genes regarding the median absolute deviation in expression profiles, which resulted in seven identified modules (Figure 4A). The *minimum module* size was 20 and the *pickSoftThreshold* was 4 (Supplemental Figure 1). The modules are referred to by their color labels in a hierarchical cluster dendrogram (Figure 4A).

**Fig. 4.**
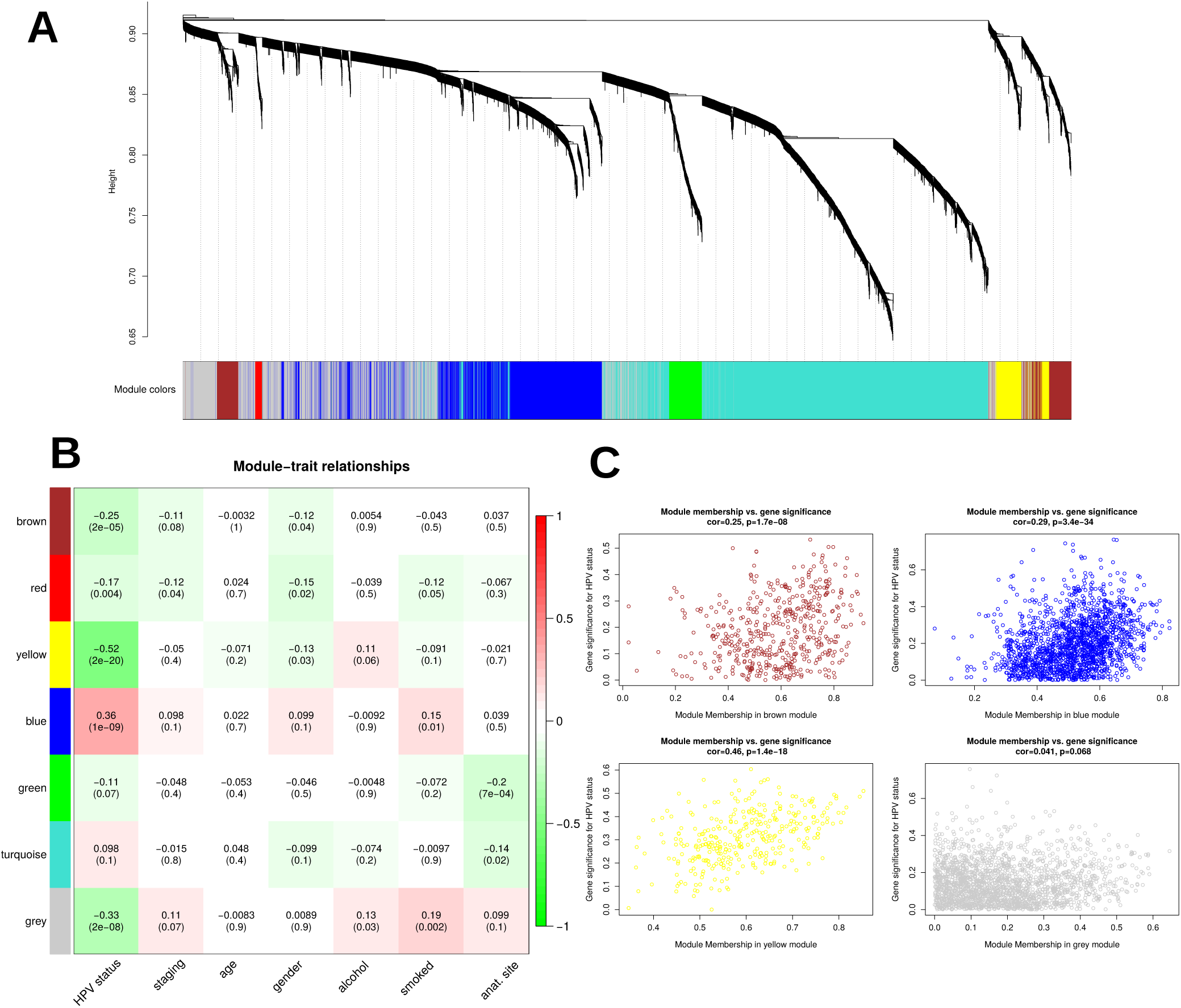
Co-expression genes modules and their relationship with studied traits. (**A**) Gene clustering dendrogram showing co-expressed genes hierarchically clustered into modules, which were defined by different colors (rectangle at bottom). (**B**) Matrix showing the correlation of the color-coded modules (rows) with studied traits (columns). Cell contents display the correlation coefficients and p-values (in parentheses). Correlation coefficients were color-coded according to the heat index from red to green depicted at the vertical bar at the right to the graph. (**C**) Scatter plots of gene significance (*y*-axis) versus module membership (*x*-axis) for HPV- status associated with selected modules with the largest significance when comparing HPV+ and HPV- samples (*Brown, Blue, Yellow and Grey*).

Figure 4B depicts the correlations of the eigengene modules with the traits including ‘HPV status’, ‘staging’, ‘age’, ‘gender’, ‘alcohol’, ‘smoked’, and ‘anatomical site’. Three modules were found significantly associated with the HPV status (absolute correlation ≥ 0.25 and p-value ≤ 0.001), *Blue, Yellow* and *Grey*. The module membership (MM) versus gene significance (GS) plot for these modules and the borderline module *Brown* are shown in Figure 4C. Despite candidate genes with no distinct module assignment were grouped in the *Grey* module, we have decided to include this module to subsequent analysis of the networks due to its significant association with HPV status. Therefore, the *Blue, Yellow* and *Grey* modules were further studied. We also computed the hierarchical clustering of the expression and methylation data of the samples concerning HPV status or disease staging using the ‘flashcluster’ function of WGCNA, but no clear clustering was observed (Supplemental Figure 2).

### The *Blue, Yellow* and *Grey* gene modules result in distinct networks according to HPV status

In general, the modules built by WGCNA contain a large number of genes when global expression data are used. As a consequence, some genes can be randomly associated with a specific phenotype. Thus, it is fundamental to identify relevant genes in the network, also known as hub genes, which are more likely to represent robust markers of specific phenotypes. There are different approaches to identify hubs in a network. In our approach, we used the previously selected DEG to guide the choice of hubs. For each selected module, we divided the samples by HPV status. We then computed the correlations in each status using all genes in each module (Spearman’s rank correlation coefficient). The genes selected in each module by HPV status were considered when these genes were DEG or when they were highly correlated with DEG. We applied a correlation threshold of ≥ 0.65 and applied a p-value threshold of ≤ 0.01 for both HPV- and HPV+ networks. In addition, we characterized the transcription factor genes (TF), doubly DEG/DMG, singly DMG and significantly mutated genes that engage in known protein-protein interactions (PPI) with present genes in each network (Figure 5). Of the 12 double DEG/DMG genes considered for analysis (see above), eight appeared in one of the three modules kept for further analysis.

The networks were differentially connected according to HPV status (Table 1). All three *Blue, Yellow* and *Grey* modules had more densely connected networks in the HPV+ compared to HPV- cases, as measured both by the number of nodes and of edges (Table 1). In the HPV+ *Blue* network, the *SYCP2* (synaptonemal complex protein 2) is much more densely connected to other genes when compared to the HPV- network (Figure 5A). The *C22orf45* gene, which showed few connections in the HPV+ *Blue* network, was not even evidenced in the HPV- counterpart, since no connections were established (Figure 5A). Concerning TF genes, there are also stronger network connections, and a higher number of TF genes involved, when the HPV+ network is compared to the HPV- counterpart. Most TF genes in the HPV+ network are connected into a single high-density cluster, which is not seen in the HPV- network. Also of note, the *YBX2* is a TF that appears only in the HPV+ network, and connects the *SYCP2* hub to that high-density TF hub. It is worth mentioning that all genes visualized in this network (including the TFs) are overexpressed in HPV+ tumors.

**Fig. 5.**
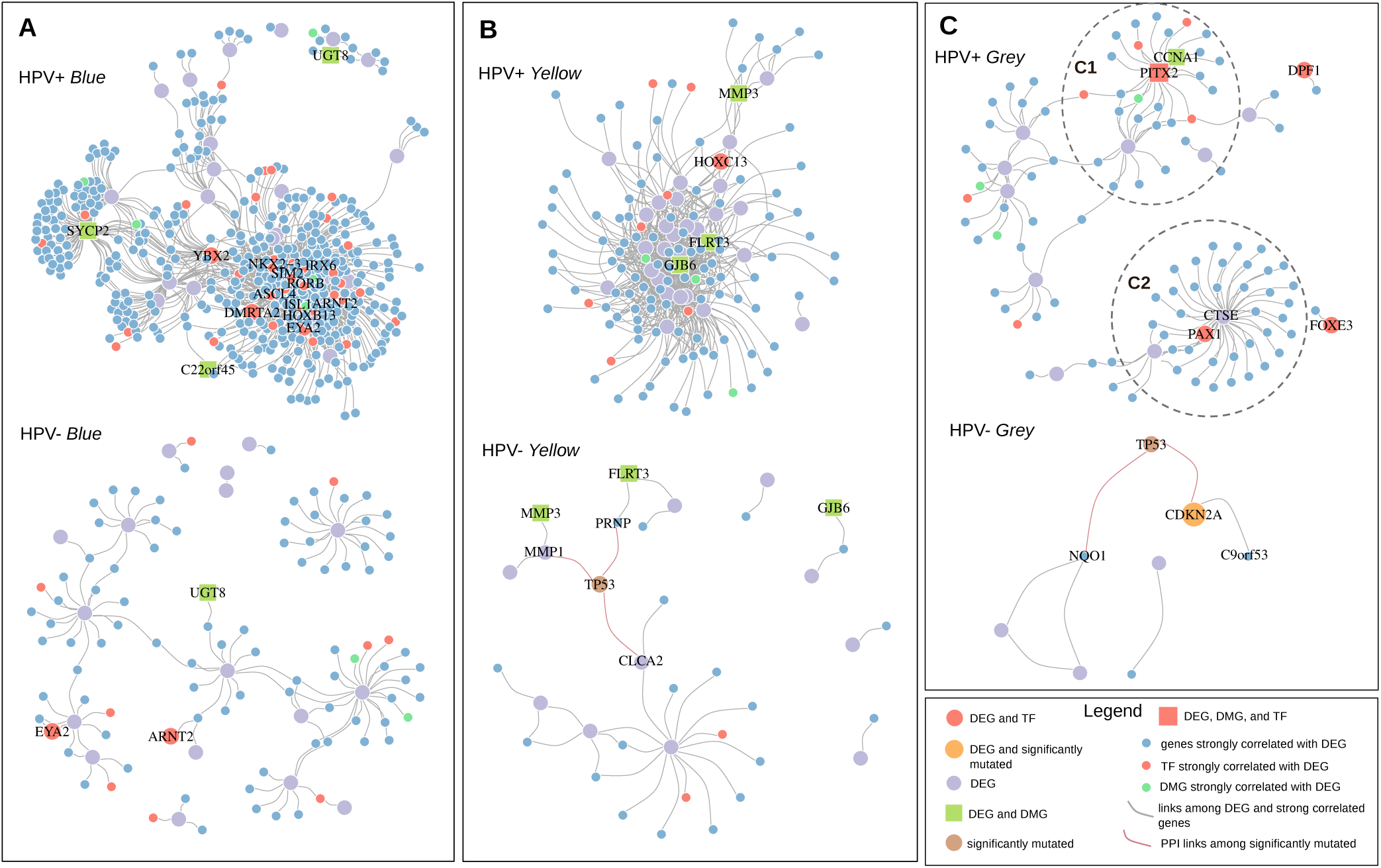
Co-expression networks among the modules with significantly different profiles between HPV+ and HPV- cases. (**A**) *Blue* module. (**B**) *Yellow* module. (**C**) *Grey* module. In all modules, gene classifications are shape- and color-coded according to the legend at the lower right inset of the Figure. Links between DEG and strongly correlated genes and also those linking significantly mutated genes with genes through protein-protein interactions are also color-coded according to the legend of the Figure.

**Table 1.** Connection metrics of co-expression networks of different modules in HPV+ and HPV- cases.

| Module | HPV+ | HPV- |
| --- | --- | --- |
| *Blue* |  |  |
| nodes | 539 | 112 |
| edges | 2633 | 114 |
| *Yellow* |  |  |
| nodes | 145 | 36 |
| edges | 640 | 34 |
| *Grey* |  |  |
| nodes | 127 | 8 |
| edges | 111 | 7 |

The *Yellow* modules (Figure 5B) depict genes that are generally overexpressed in HPV- compared to HPV+ tumors. In this set, *MMP3, FLRT3* and *GJB6* were doubly selected (in expression and methylation analyses) and more tightly connected in HPV+ tumors, denoting a concerted downregulated pathway. The *HOXC13* TF is also underexpressed in HPV+ tumors, and likely plays an important role in the connexion of the pathways encompassing those genes.

The *Grey* modules (Figure 5C) encompass genes that were not consistently clustered into any of the modules characterizing definite co-expression profiles. However, in the HPV+ network, specific co-expression gene sub-networks can be retrieved that show under or overexpression compared to HPV- tumors. Genes placed in central hubs of these two subnetworks can be visualized in the C1 and C2 inset circles of Figure 5C, respectively. TFs which are DEG and/or DMG and involved in the control of these sub-networks include *PITX2* in C1 (underexpressed in HPV+ tumors) and *PAX1* in C2 (overexpressed in those tumors).

### Significantly mutated genes are evidenced only in HPV- HNSCC

The top 10 frequently mutated genes derived from our studied dataset were analyzed considering samples independently of their disease stage classification and HPV status. A comprehensive annotation of all the types of mutations found for those top 10 mutated genes is shown in the detail in Figure 6A. As expected and previously described in the original TCGA study (10), the TP53 gene was the most frequently mutated, followed by *FAT1* and *CDKN2A*, all highly mutated in HPV- tumors (82%, 27% and 25% of those tumors, respectively). On the other hand, *PIK3CA, CSMD3* and *HUWE1* were the most frequently mutated genes found among the HPV+ samples (Figure 6A and Supplemental Figure 3). We next applied a Fisher’s exact test in all dataset to obtain the differentially mutated genes between the HPV+ and HPV- statuses with an adjusted p-value < 0.05 provided by the FDR estimate. Only the top three genes, *TP53, FAT1* and *CDKN2A,* were significantly mutated in the HPV- cases, whereas no significantly mutated gene was found in the HPV+ cases (Figure 6B).

**Fig. 6.**
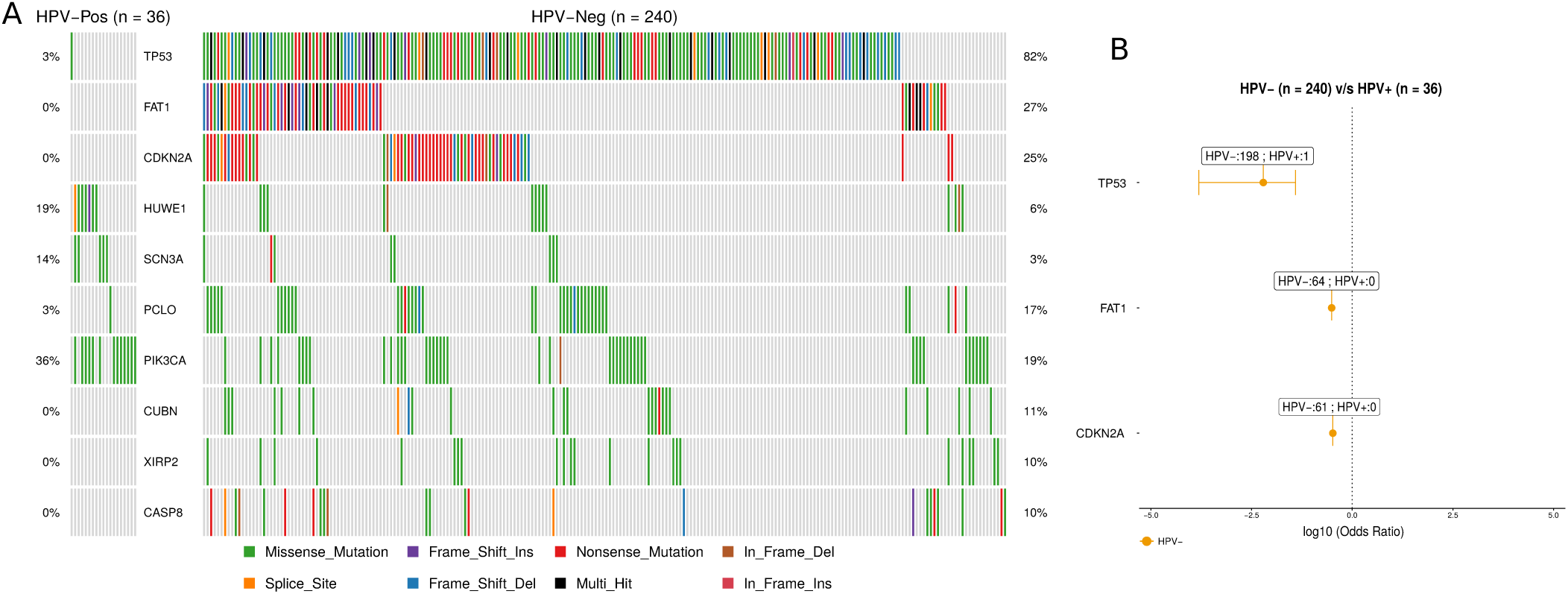
Mutation burden in HPV+ and HPV- HNSCC samples. (**A**) Top 10 frequently mutated genes in HPV+ (n = 36) and HPV- (n = 240) HNSCC cases, sorted according to their frequency in the latter. Each case is represented by a single column in both the HPV+ and HPV- graphs. Relative mutation frequencies are shown at the left of the HPV+ graph and at the right of the HPV- graph. Types of mutations found in each gene were color-coded according to the legend at the bottom of the Figure. (**B**) Forest plot showing the odds ratio (OR) of the significantly mutated genes using the Fisher’s exact test (FDR-adjusted p-value < 0.05). Only three genes (*TP53, FAT1* and *CDKN2A*) were retrieved with significance, all found in HPV- cases. Values are plotted in a log10 scale.

The three abovementioned significantly mutated genes had their locations and relationships visualized in the HPV- networks shown in Figures 5A-C (bottom panels). *TP53* appears in two of the HPV- modules (*Yellow* and *Grey*), while *CDKN2A* appeared only in the *Grey* module, as it is also a DEG in that case. In the HPV- *Yellow* module, *TP53* appears connected with *MMP1, CLCA2* and *PRNP* (Figure 5B, bottom panel). In the *Grey* module, *TP53* evidences a connection with *CDKN2A* and with *NKO1*, while *CDKN2A* itself is additionally associated with *C9orf53* (Figure 5C, bottom panel).

### Enrichment functional analysis highlights specific HPV+ and HPV- biological pathways

To further explore the possible role of the gene modules and networks identified in our analyses of HNSCC with distinct HPV statuses, we performed enrichment analysis with Gene Ontology (GO) - Biological Process (BP), the Kyoto Encyclopedia of Genes and Genomes (KEGG) and the Molecular Signatures Database (MSigDB). The hypergeometric analysis was conducted with Bonferroni adjusted FDR ≤ 0.05. We captured the enriched functions of the identified modules with the R ‘clusterProfiler’ package. The main results are shown in Table 2. We identified pathways associated with cell fate specification and glycolysis/gluconeogenesis in the Blue HPV+ module. In contrast, genes of the HPV+ *Yellow* module were downregulated in the overrepresented biological processes of epidermis development, negative regulation of epithelial cell proliferation and keratinocyte differentiation. Finally, processes and pathways involving dendritic spine morphogenesis and lysosome degradation were overrepresented in HPV+ tumors, while cellular ketone metabolism and aging were underpinned in HPV- tumors (Table 2).

**Table 2.**
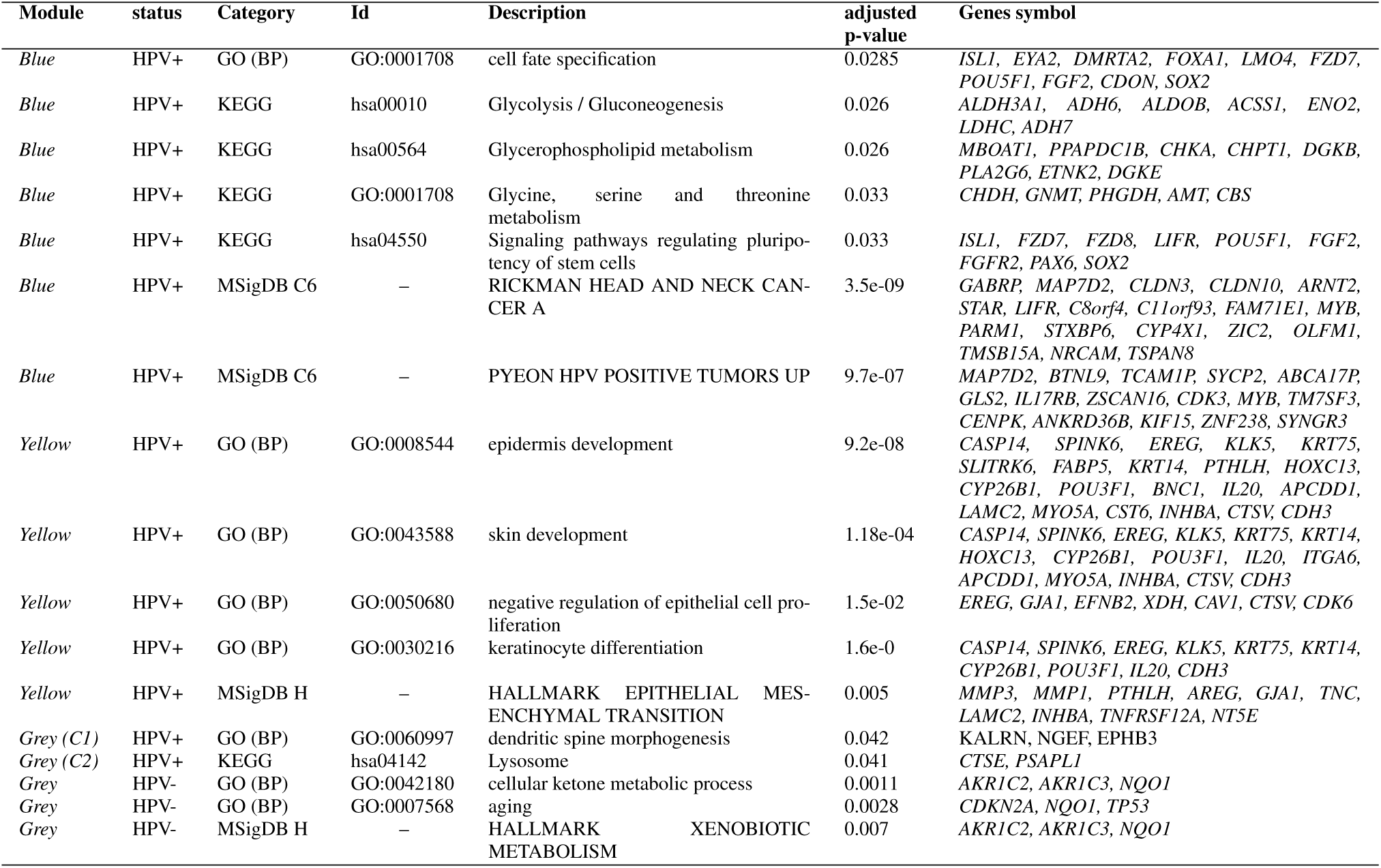
Biological processes, pathways and molecular signatures significantly overrepresented in the Blue, Yellow and Grey modules according to HPV status.

## Discussion

The heterogeneity of HNSCC with respect to the variety of anatomical sites and driving behavioral (alcohol, tobacco, hot beverages) and infectious (HPV) factors makes the identification of relevant therapeutic targets a challenging task (9, 28, 29). In addition, the analysis of mono-omic data, i.e. from a single layer, provides only one dimension of a multifaceted scenario, and limited information about the possible molecular mechanisms involved in the disease. Genetic and epigenetic changes such as mutations and methylation patterns modulate gene expression levels of several genes. Although both result in the same phenotypic alteration (changes in expression levels), the genetic mechanisms involved and the adjacent gene interactions (gene networks) are different, an observation that can only be done with the analysis of multiomic data (21, 30, 31). In this sense, the analysis of data such as those available through the TCGA Consortium provides a unique opportunity to assess multi-layer molecular interactions in a feasible manner (24, 32). In the current study, we utilized HNSCC multi-omic data from TCGA in an attempt to more comprehensively understand gene co-expression networks and the putative roles of gene promoter methylation patterns and gene mutations associated with HPV+ and HPV- profiles through disease progression.

Our approach started with the identification of DEG between HPV+ and HPV- tumors. The genes involved in these networks appeared to vary significantly when we use data from different disease stages (I through IV) of both HPV statuses, indicating that their expression is modulated to different levels during the carcinogenic process. Despite the fact that most genes lost statistical significance in differential expression between HPV+ and HPV- tumors at one or more disease stages, a general trend could be observed that DEG maintained their patterns throughout the stages (i.e., being over or underexpressed in HPV+ compared to HPV- cases). It is worth mentioning that data from a small number of cases were available for initial tumor stages, particularly from HPV+ cases, and we cannot exclude the possibility that such heterogeneity in the number of samples compromised the robustness of the differences observed herein. Analyses with larger numbers of cases are warranted in further studies to more precisely identify DEG throughout HNSCC stages. On the other hand, most of the converging expression results (i.e., lack of significant differential expression between HPV+ and HPV- cases) occurred in stage IV (data not shown). It is tempting to speculate that, at an advanced disease stage, the molecular processes converge between HPV+ and HPV- cases, being the virus a mere initiator of the carcinogenesis via distinct pathways. A similar scenario was observed when DMG were derived from the same data. Again, promoter gene methylation patterns differed between HNSCC stages comparing HPV+ and HPV- cases and no single gene differed significantly across all four stages between the two HPV statuses. These results indicate that the association between methylation and gene expression is stronger in HPV-infected HNSCC, and thus that epigenetic regulation appears to be pivotal during HPV infection of head-and-neck anatomical sites.

Gene co-expression modules and networks were constructed using global expression data, and DEG were used as filters for refining those networks as described in Methods. In this sense, only genes that were DEG or directly interact with DEG in a linear positive fashion were plotted in module networks. Three significantly supported modules (*Blue, Yellow* and *Grey*) were further investigated. The obtained gene networks differed between HPV+ and HPV- tumors within each module, suggesting that HPV infection plays a unique role in HNSCC carcinogenesis, which involves a series of distinct molecular processes from the HPV- counterparts. To the best of our knowledge, very few studies (if any) tried to assess the composition of gene networks through disease progression and also how HPV influences that development. In all three modules studied, the HPV+ networks were much more densely connected and encompassed a larger number of significant nodes and edges compared to the HPV- counterparts. Irrespective of the modulation provided by the presence of the virus (either gene overexpression or underexpression compared to the HPV- networks in the *Blue* or *Yellow* module, respectively), the networks suggest a fundamental role of HPV in hijacking and modulating specific biological processes within tumor cells.

We further integrated DMG and significantly mutated genes in the HPV+ and HPV- module networks by identifying those genes within the networks. Since the networks are composed of DEG, the DMG identified are necessarily doubly DEG and DMG. Despite these genes were essentially the same when comparing the HPV+ and HPV- networks within each module (with one or two exceptions in each module), their engagement in different interactions were strongly dispar. In the HPV+ networks, particularly in the *Yellow* module, DMG were more central in the networks and engaged in higher numbers of connections. This is consistent with the observation that DEG in the *Yellow* module are repressed (underexpressed) in the HPV+ cases compared to HPV- counterparts. With respect to the mutated genes, three appeared significantly mutated in HPV- compared to HPV+ cases, *TP53, CDKN2A* and *FAT1*. Of those three, only *CDKN2A* appeared in one of the modules (*Grey*), because it is also a DEG. On the other hand, *TP53* also showed relevant PPI with genes in the *Yellow* and *Grey* modules as evidenced through searches within StringDB, as was arbitrarily added to those two modules. As expected, all those occurrences took place in the HPV- networks, consistent with the fact that mutations in those genes were reported almost exclusively in subjects with HPV- status (see Figure 6B). Our results point to a fundamental, yet expected role of host gene mutations as primary drivers of carcinogenesis in HPV- samples, as opposed to an infectious agent driver in the case of HPV+ samples. Of note, mutations in the *PIK3CA* (phosphatidylinositol-4,5-bisphosphate 3-kinase catalytic subunit alpha) gene have been recently associated with HPV+ HNSCC (10, 33). We did not find such association, since those mutations were also present in HPV- tumors, and the difference between the two HPV statuses was not significant (19% of HPV- versus 36% of HPV+ cases), a comparison likely not conducted by Lawrence et al. in their report (10).

Several genes could be retrieved from the gene co-expression networks obtained from the three modules which appear to have distinguishable importance in HPV+ and HPV- HNSCC. The *SYCP2* gene encodes the synaptonemal complex protein 2, a protein that is localized in chromosomal centromeres and responsible for the association of chromosomes with the synaptonemal complex, driving the prophase of meiosis (34). SYCP2 has been found overexpressed in HPV+ oropharyngeal cancers (11), and similar results were found herein for HNSCC in general. It was also associated with cervical squamous cell carcinomas (35). Of note, Peyon et al. have proposed that HPV+ cancers from distinct anatomical sites, specifically cervical cancers and HNSCC, share many upregulated genes and pathways, including the overexpression of testis-specific genes involved in meiosis such as *SYCP2* (36). Aberrant expression of this gene in HPV+ cancers likely contribute to genomic instability and further oncogenic alterations, yet a specific interaction of viral products with *SYCP2* is yet to be elucidated.

Transcription factors were also overexpressed in the HPV+ *Blue* network, such as *YBX2, DMRTA2* and *EYA2*. Most of these TF have been described as overexpressed in different types of cancers, including ovarian, testis and breast cancer in the case of *YBX2* (37, 38), and breast cancer, lung cancer and acute myeloid leukemia in the case of *EYA2* (39, 40). *DMRTA2*, on the other hand, is also expressed in the spermatogenesis of the testis, and regulates the cyclin-dependent kinase CDKN2c (41), in addition to maintaining neuroprogenitor cells in the cell cycle (42). Although the specific role of HPV in upregulating these TF is unknown, gene silencing of *EYA2* significantly reduced viability, migratory capacity, and anchorage-independent growth of HPV16-transformed keratinocytes (43). Moreover, our results point to a fundamental interaction of HPV with a defined network of genes that regulate gametogenesis in the testis and ovaries, a pathway that warrants further study for interventional approaches. Additional genes that are co-expressed with the abovementioned ones in a highly significant fashion, such as *MYO3A* (myosin IIIA), *IL17RB* (interleukin 17 receptor B) and *UBXN11* (UBX domain protein 11), and for which scarce information as related to carcinogenesis or HPV infection is available, are also attractive for further studies and as targets for intervention. According to the GO biological processes associated with the reconstructed HPV+ *Blue* network, cell fate differentiation and glucose metabolism appear to be major components, consistent with gene upregulation that occurs during tumor development.

In the HPV+ *Yellow* network, two central genes were shown to be significantly underexpressed and more methylated compared to HPV- HNSCC, *GJB6* (gap junction protein beta 6, also known as connexin 30) and *FLRT3* (fibronectin leucine rich transmembrane protein 3). Furthermore, the *HOXC13* (homeobox C13) TF, a regulator of several genes during epithelial differentiation, and of which mutations were associated with pure hair and nail ectodermal dysplasia (44), is also underexpressed in this HPV status. Conversely, *HOXC13* and *FLRT3,* among other genes seen in our *Yellow* networks, were found upregulated in HPV- OSCC (45), in agreement with our results. Not surprisingly, all these genes have been associated with the expression and metabolism of gap junction proteins and keratins, as well as keratinocyte differentiation in epithelial cells, and appeared to be downregulated in HPV+ tumors. Other genes significantly associated with those are keratins 14 and 19 *(KRT14, KRT19), COL4A6* (collagen type IV alpha 6 chain) and *CLCA2* (chloride channel accessory 2), which are also involved in keratinocyte biology. These results are in consonance with the GO analysis for this network, which showed an enrichment in negative regulation of epithelial cell proliferation, keratinocyte differentiation, and skin and epidermis development (Table 2). The *MMP3* gene encodes the matrix metallopeptidase 3 and is generally associated with multiple steps of cancer development, invasion and metastasis (46). Interestingly, this gene was also underexpressed in our HPV+ compared to the HPV- *Yellow* network. It is tempting to speculate that, in a scenario where most adhesion and gap junction molecules are already downregulated, upregulation of *MMP3* is not a *sine qua non* step towards tumor cell invasion and metastasis.

In the HPV+ *Grey* network, two genes were found underexpressed and hypermethylated compared to HPV- tumors, *PITX2* (paired like homeodomain 2) and *CCNA1* (cyclin A1). The first one is additionally a TF which has been implicated in muscle development. *PITX2* hypermethylation has been interestingly associated with better prognosis in HNSCC (47) but with worse prognosis in breast cancer (48). *PITX2* has also been shown to control the expression of *CCNA1* in a positive fashion (49), which fits the relationships found in our network. Moreover, HPV-16 E7 has also been implicated in the mediation of *CCNA1* promoter methylation (50). Conversely, *PAX1* (paired box 1) and correlated genes (Figure 5C, inset C2) are overexpressed in HPV+ compared to HPV- tumors. One of these genes, the DEG *CTSE* (cathepsin E), is involved in the lysosome degradation pathway (KEGG, hsa04142). *CTSE* has been additionally recognized as a biomarker for the detection of pancreatic ductal adenocarcinoma (51) and for gastric cancer (52).

In the HPV- *Grey* module, no clear networks were formed, but underexpression of *CDKN2A* and its association with *TP53* were evident. Moreover, *TP53* and *CDKN2A* were significantly mutated in this network. *CDKN2A* is a kinase implicated in the production of p16(INK4a) and p14(ARF), well-established tumor suppressors. Therefore, decreased expression of *TP53* and *CDKN2A* by inactivating mutations as seen in our data fits the scenario of HPV- induced carcinogenesis, where cellular genes are the major drivers of the process.

Overall, the results presented herein emphasize the importance of integrating different genomic data (as mRNA expression, DNA methylation and mutation patterns) to get a better understanding of the molecular mechanisms involved in the carcinogenesis and progression of HNSCC, an approach that can be applied to other tumor types. Even though the individual analysis of one biological level (mRNA) gives information associated with the disease, the integration with other biological levels is required to have a more comprehensive view from a functional perspective, allowing the identification of novel molecular targets unseen by mono-omic approaches.

## Methods

### Omics datasets and preprocessing

The multi-omic data of HNSCC were retrieved from The Cancer Genome Atlas (TCGA) database (24) by selecting the datasets published in 2015 (10) which identified HPV-positive (HPV+) and HPV- negative (HPV-) cases, totalling a set of 279 patient with data of primary solid tumors. Using clinical data information, we grouped the samples by HNSCC staging, which excluded three patients for whom this information was absent. The resulting dataset for further analysis consisted of 240 HPV- and 36 HPV+ cases.

The gene expression dataset was composed of data generated in an Illumina HiSeq 2000 RNA-Seq platform (level 3) using the preprocessed RNAseqV2 normalized count expression values based on RNA-Seq by Expectation-Maximization (RSEM). We performed a log-transformation *log*(1 + *p*) on the count expression values. Genes with a zero standard deviation were removed from the dataset.

The methylation dataset was determined using Infinium HumanMethylation450 BeadChip array. In the methylation level 3 data, each probe (CpG site) is measured as the ratio (*β* value) of the signal of methylated probes with respect to the sum of methylated and unmethylated probes, which varied continuously from 0 to 1, values that indicate *unmethylated* and *fully methylated,* respectively. We removed cross-reactive, non-specific, single nucleotide polymorphisms (SNPs), chromosomes *X* and *Y* and probes with genomic coordinates set to zero. We also removed probes with more than 5% missing values across samples. In the remaining data, absent data were estimated using the weighted k-nearest neighbor (kNN) algorithm, with k = 10, as proposed by Troyanskaya (2001) (53) and implemented in the R ‘impute’ package. The raw data (*M* values) normalization was performed with the BMIQ method proposed by Teschendorff (2013) (54) and implemented in the Chip Analysis Methylation Pipeline (ChAMP) (55). The analysis of DMG was performed with the defined promoter region, following the methodology used by Jiao (2014) (56). Briefly, the average value of the probes mapping within 200 bp of the transcription start site (TSS) was assigned to the gene. If no probes mapped within 200 bp of the TSS, we used the average value of probes mapping to the 1st exon of the gene. If such probes were also not available, we used the average value of probes mapping within 1500 bp of the TSS.

The somatic mutation data were obtained from the Mutation Annotation Format (MAF) files. MAF files provide baseline data for many downstream analyses identifying somatic mutations in cancers through various pipelines and sequencing platforms. MAF files provide baseline data for many downstream analyses identifying somatic mutations in cancers through various pipelines and sequencing platforms.

### Genes selected by differences among stages in expression and methylation data

We selected significant genes (False Discovery Rate, FDR-adjusted < 0.01) comparing each profile (HPV+ versus HPV-) for each HNSCC stage. For instance, HPV+ (stage I) vs HPV- (stage I), …, HPV+ (stage IV) vs HPV- (stage IV). Genes that were selected in at least one comparison were included in posterior analyses. We used this approach for the RNA-Seq dataset including absolute Fold-Change (absFC) ≥ 4, resulting in Differentially Expressed Genes (DEG). For the methylation dataset, we used the same method but considering the absFC ≥ 5 for selecting the DMG. These analyses were achieved based on normalized datasets by the fitting of the linear model (for each probe or gene) followed by moderated t-tests implemented in the ‘limma’ package (25). We overlapped the DEG and DMG to determine genes that were doubly selected. Next, we calculated the Pearson’s correlation (PC) between the methylation and expression values to those doubly selected genes.

### Somatic mutation analysis

Somatic mutations from Whole Exome Sequencing (WXS) in HNSCC were downloaded in a MAF file. We performed Fisher’s exact test to detect the differentially mutated genes on all HPV+ versus HPV- profiles with the ‘maftools’ package (57). We used the adjusted p-value ≤ 0.05, given by the FDR analysis (58).

### Co-expression modules via WGCNA

The analysis of the co-expression network modules was performed using the package Weighted Correlation Network Analysis (WGCNA) (59), applying the *minimumModuleSize* = 20 and *mergingCutHeight* = 0.45. The similarity matrix was converted to a weighted adjacency matrix by raising it to the power of *β* to amplify the strong connections and penalize the weaker connections. The trait-associated mRNAs are then subjected to WGCNA (60) for the identification of high coexpression modules, denoted as *M*. The clinical data used in the analysis was related with ‘HPV status’, ‘staging’, ‘age’, ‘gender’, ‘alcohol’, ‘smoked’, and ‘anat. site’. A subset *M’* of *M* is given by modules significantly associated with HPV status selected for posterior analysis (absolute correlation > 0.25 and p-value < 0.001).

### Refining modules and interactions networks

Due to the number of genes in high-throughput data, the resulting modules contain a large number of genes, with interconnections that might result from spurious correlations. In order to obtain a selective and restrictive set of genes involved in each profile, we filtered the nodes in HNSCC for HPV+ and HPV- phenotypes. For this, assuming we have *n* selected modules, each selected module 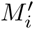 of *M′* ⊆ *M* is represented by

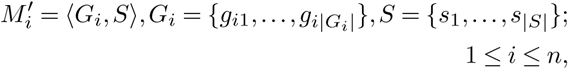

where *G_i_* is a set of genes and *S* is the set of samples. We separated the modules in,

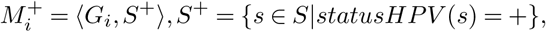

for HPV+ and

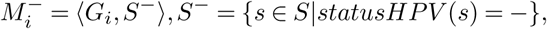

for HPV- . For each 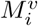, 0 ≤ *i* ≤ *n* where *υ* ∈ {+, -},

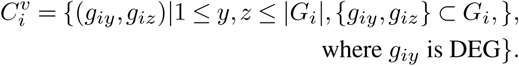

We select the genes *g_iy_* which *corr_S_^υ^* (*g_iy_, g_iz_*) ≥ 0.65, *y* > *z*, (*g_iy_, g_iz_*) ∈ 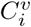 and p-value < 0.01.

The resulting networks were visualized with the ‘igraph’ package available in R CRAN (61). DEG, DMG, doubly selected (DEG and DMG), transcription factors (TF) and significantly mutated genes were identified in the network. The TF data were obtained from the *TFcheckpoint* database (62). To link the significantly mutated genes in the network, we used the protein-protein Interaction (PPI) associations from the STRING database (63), with high confidence (score ≥ 0.7) selected.

### Gene Ontology and pathway-enrichment of the selected genes within modules

To identify the significant enrichment pathways, Gene Ontology (GO) terms (64, 65), KEGG (66) and the Molecular Signature Database (MSigDb v.6.0) (67) were used. The hypergeometric distribution test (p-value < 0.001) was used to test for statistically significant overrepresentation of genes from particular biological gene sets within the co-expression in each module and HPV status. The p-values were corrected for multiple testing through FDR using the R package ClusterProfiler (68).

## Availability of supporting data and materials

The networks generated in the analysis above are available in an interactive module at: https://quelopes.github.io/files/projects/HNSCC/Co-expressionHNSCC.html. The HPV+ networks were modeled and populated in the graph database Neo4J. The database can be retrieved at GitHub <https://github.com/quelopes/HNSCC-network>.

## Additional files

### Additional Figure 1

(Additional_file_1.pdf): Hierarchical clustering of the expression (top panel) and methylation (bottom panel) data of the samples with respect to HPV status or disease staging (horizontal bars at the bottom of each dendogram, color-labeled according to the legend at the bottom of the Figure.

### Additional Figure 2

(Additional_file_2.pdf): Scatter plot of network topology analysis using a range of soft-thresholding powers. Top left, scale-free fit index as a function of the soft-thresholding power indicating the soft-thresholding power of 4. The other charts represent the mean connectivity (top right), centralization (bottom left) and heterogeneity (bottom right) (*y*-axes) *versus* soft-thresholding power (*x*-axes).

### Additional Figure 3

(Additional_file_3.pdf): Mutation patterns in HPV+ (right panel) and HPV- (left panel) samples. For each HPV status, the plots show variant types, variant classifications, single nucleotide variation (SNV) classes, variants per sample, variant classification summary and frequently mutated genes according to variant classification.

## Abbreviations

ChAMP: Chip Analysis Methylation Pipeline
CRAN: Com-prehensive R Archive Network
DEG: Differentially Ex-pressed Genes
DMG: Differentially Methylated Genes
FDR: False Discovery Rate
GS: gene significance
HNSCC: Head-and-Neck Squamous Cell Carcinoma
HPV: Human papillomavirus
KNN: k-Nearest Neighbor
MAF: Mutation Annotation Format
MM: module membership
OPSCC: Oropharyngeal Squamous Cell Carcinoma
PC: Pearson’s correlation
PPI: Protein-Protein Interaction
SNP: Single Nucleotide Polymorphism
TCGA: The Cancer Genome Atlas
TIL: tumor-infiltrating lymphocytes
TF: Transcription Factor
TSS: Transcription Start Site
WGCNA: Weighted Gene Co-expression Network Analysis
WXS: Whole Ex-ome Sequencing.

## Ethical Approval (optional)

Not applicable.

## Consent for publication

Not applicable.

## Author’s Contributions

RLC, MB and MAS conceived the study. RLC and MB designed the experiments. RLC analyzed the data and prepared figures and tables. All authors wrote the manuscript, reviewed its drafts, approved its final version and agreed with its submission.

## Acknowledgements

We would like to thank the Brazilian National Laboratory of Scientific Computing (LNCC), Brazilian Ministry of Science and Technology, for providing computational infrastructure to analyze the data of the study. We would also like to thank Dr. Nicole Scherer for providing additional support to the use of the infrastructure from the Bioinformatics and Computational Biology Lab of INCA, Brazilian Ministry of Health.

